# Identification of a Novel Neutralizing and Two Non-Neutralizing Epitopes on Epstein-Barr Virus gp350 Protein

**DOI:** 10.1101/302844

**Authors:** Lorraine Z Mutsvunguma, Anne Barasa, Charles Warden, Joslyn Foley, Murali Muniraju, David H Mulama, Peng Ye, Hanjun Qin, Jinhui Wang, Xiwei Wu, Waithaka Mwangi, Javier Gordon Ogembo

**Affiliations:** Department of Immuno-Oncology, Beckman Research Institute of City of Hope, Duarte, California, USA; Department of Human Pathology, University of Nairobi, Nairobi, Kenya; Integrative Genomics Core, Beckman Research Institute of City of Hope, Duarte, California, USA; Department of Biological Sciences, Masinde Muliro University of Science and Technology, Kakamega, Kenya; Department of Diagnostic Medicine/Pathobiology, College of Veterinary Medicine, Kansas State University, Manhattan, Kansas, USA

**Keywords:** EBV, glycoprotein, gp350, antibodies, epitopes, amino acid residues, vaccines

## Abstract

Prevention of Epstein-Barr virus (EBV) primary infection has focused on generating neutralizing antibodies (nAbs) targeting the major envelope glycoprotein gp350/220 (gp350). To date, eight gp350 epitopes have been identified, but only one has elicited nAbs. In this study, we generated 23 hybridomas that produced anti-gp350 antibodies. We compared the candidate anti-gp350 antibodies to nAb-72A1 by: (1) testing their ability to detect gp350 using ELISA, flow cytometry, and immunoblot; (2) sequencing their heavy and light chain complementarity-determining regions (CDRs); (3) measuring the ability of each monoclonal antibody (mAb) to neutralize EBV infection *in vitro*; and (4) mapping the gp350 amino acids bound by the mAbs using RepliTope peptide microarrays. Eight antibodies recognized both denatured and non-denatured gp350, whereas five failed to react with denatured gp350 but recognized native gp350, suggesting they recognized conformational epitope(s). Sequence analysis of the heavy and light chain variable regions of the hybridomas identified 15 as mAbs with novel CDR regions unique from those of nAb-72A1. Seven of the new mAbs neutralized EBV *in vitro*, with HB20 and HB17 reducing EBV infection by 40% and >60%, and >30% and 80%, at 10 μg/ml and 50 μg/ml, respectively. Epitope mapping identified nine epitopes and defined their core residues, including two unique immunodominant epitopes, _253_**TPIPGTGYAYSLRLTPRPVSRFL**_253_ and _875_**LLLLVMADCAFRRNLSTSHTYTTPPY**_899_, and a novel nAb epitope _381_**GAFASNRTFDIT**_392_. This study provides comprehensive *in vitro* mapping of the exact residues defining nine epitopes of EBV gp350. Our findings will inform novel strategies to design optimal EBV vaccines capable of conferring broader protection against the virus.

**Importance:** Neutralizing antibodies (nAbs) directed against Epstein-Barr virus envelope glycoprotein gp350/220 (gp350) are generated in humans upon infection or immunization, and are thought to prevent neonatal infection. However, clinical use of exogenous nAbs (passive immunization) is limited to a single study using the only well-characterized nAb, 72A1. The gp350 ectodomain contains at least eight unique B-cell binding epitopes; two of these epitopes are recognized by nAb-72A1. The exact amino acid residues of the other six epitopes and their role in generating nAbs has not been elucidated. We used our 15 newly generated and fully characterized monoclonal antibodies and a peptide-overlapping RepliTope array to provide a comprehensive map of the core amino acid residues that define epitopes of gp350 and to understand their role in generating nAbs. These results will inform design of better-targeted gp350 peptide vaccines that contain only protective epitopes, which will focus the B-cell response to produce predominantly nAbs.

## Introduction

Epstein-Barr virus (EBV) infection is the causal agent of acute infectious mononucleosis (9, 13). Persistent EBV infection in immunodeficient individuals is associated with numerous epithelial and lymphoid malignancies, such as nasopharyngeal carcinoma, gastric carcinoma, Burkitt lymphoma, Hodgkin lymphoma, and post-transplant lymphoproliferative diseases (PTLD) (24). Pre-existing antibodies provide the primary defense against viral infection. Prophylactic prevention of EBV primary infection has mainly focused on blocking the first step of viral entry by generating neutralizing antibodies (nAbs) that target EBV envelope glycoproteins. Five glycoproteins in particular—gp350/220 (gp350), gp42, gH, gL, and gB—are required for efficient infection of permissible host cells and have emerged as potential prophylactic targets (2, 3, 5, 20).

EBV predominantly infects epithelial cells and B cells, reflecting the viral tropism and the cellular ontogeny for EBV-associated malignancies (4). There are two schools of thought on how the initial EBV transmission into the human host cells occurs. In the first infection model, the incoming virus engages with ephrin receptor A2 via heterodimeric gH/gL, which triggers gB fusion with the epithelial cell membrane and entry of the virus into the cytoplasm (4). This interaction is thought to occur in the oral mucosa, where the virus undergoes lytic replication to release virions that subsequently infect B cells. In the alternative model, the incoming virus binds to the host cell via complement receptor type 1 (CR1)/CD35 (17) and/or CR2/CD21 through its major immunodominant glycoprotein, gp350 (6). The interaction between gp350 and CD35 and/or CD21 triggers viral adsorption, capping, and endocytosis into the B cell (31), which subsequently leads to the heterotrimeric viral glycoproteins complex, gp42/gH/gL, binding to HLA class II molecules to activate gB membrane fusion and entry. Because these two models are not necessarily mutually exclusive, and given that both gp350 and gH/gL complex are important in initiating the first viral contact with host cells, use of nAbs that target either gp350 or gH/gL complex, or both, may potently block incoming virus at the oral mucosa.

Nearly all EBV-infected individuals develop nAbs directed to the ectodomains of these glycoproteins (25, 36). These antibodies can prevent neonatal infection, can protect against acute infectious mononucleosis in adolescents, and can protect against several human lymphoid and epithelial malignancies associated with EBV infection (7, 14, 23, 28). Although numerous monoclonal antibodies (mAbs) have been generated against EBV gp350 (11, 21, 34), only two murine mAbs, the non-neutralizing 2L10 and the neutralizing 72A1, have been extensively characterized and made commercially available (11, 34). Importantly, nAb-72A1 conferred shortterm clinical protection against EBV transmission after transplantation in pediatric patients in a small phase I clinical trial (8).

EBV gp350 is the most immunogenic envelope glycoproteins on the virion. It is a type 1 membrane protein that encodes for 907 amino acid (aa) residues. A single splice of the primary transcript deletes 197 codons and joins gp350 codons 501 and 699, in frame, to generate the gp220 messenger RNA. Both gp350 and gp220 are comprised of the same 18-aa residue at the C terminus that is located within the viral membrane, a 25-aa residue at the transmembrane-spanning domain, and a large highly glycosylated N-terminal ectodomain, aa 1–841 (32). The first 470 aa of gp350 are sufficient for binding CD21 in B cells, as demonstrated by a truncated gp350 (aa 1–470) blocking the binding of EBV to B cells and reducing viral infectivity (8). The gp350-binding domain on CD21 maps to N-terminal short consensus repeats (SCRs) 1 and 2, which also bind to a bioactive fragment of complement protein 3 (C3d) (16, 18). A soluble truncated EBV gp350 fragment (aa 1–470) and soluble CD21 SCR1 and SCR2 can block EBV infection and immortalization of primary B cells (32). However, gp350 binding to CD35 is not restricted to N-terminal SCRs; it binds long homologous repeat regions as well as SCRs 29–30 (22).

The gp350 ectodomain is heavily glycosylated, with both N- and O-linked sugars, which accounts for over half of the molecular mass of the protein. Currently there is only one crystal structure available for gp350, comprised of a truncated structure between 4–443 aa, with at least 14 glycosylated arginines coating the protein with sugars, with the exception of a single glycan-free patch (30). Mutational studies of several residues in the glycan-free patch resulted in the loss of CD21 binding (30), suggesting that binding of CD35 and CD21 by gp350 is mediated within this region.

There are at least eight unique CD21 binding epitopes located at the N-terminus of the gp350 ectodomain (35); at least one of these epitopes (aa 142–161) is capable of eliciting nAbs (32, 35). The aa residues 142–161 are also the binding site for nAb-72A1 (11, 30). Using gp350 synthetic peptides binding to CD21 on the surface of a B cell line, an additional gp350 epitope was identified in the C-terminal region of gp350 (aa 822–841), suggesting it is involved in EBV invasion of B cells (35). The role of other epitopes in eliciting nAbs has not been fully investigated. Furthermore, the exact aa residues that comprise the core binding sites for epitopes capable of eliciting neutralizing and non-nAbs have not been determined. Mapping the EBV gp350 protein residues that define immunodominant epitopes, identifying the critical aa residues of the known and unknown epitopes, and defining their roles in generating neutralizing and non-nAbs will guide rational design and construction of an efficacious EBV gp350-based vaccine that would focus B-cell responses to the protective epitopes.

In this study, we generated 23 hybridomas producing antibodies against EBV gp350. To assess their clinical potential and utility in informing future prophylactic and therapeutic vaccine design, we: (1) tested the ability of the antibodies produced by the new hybridomas to detect gp350 protein by enzyme-linked immunosorbent assay (ELISA), flow cytometry, and immunoblot; (2) sequenced the unique complementarity-determining regions (CDRs) of the heavy and light chains of all 23 hybridomas to identify novel mAbs; (3) measured the efficacy of each mAb to neutralize EBV infection *in vitro*; and (4) used RepliTope peptide microarrays to identify gp350 core aa residues recognized by neutralizing and non-neutralizing mAbs. Using the newly generated antibodies, we identified a new epitope bound preferentially by nAbs, distinct from the canonical neutralizing epitope bound by nAb-72A1, as well as two immunodominant epitopes bound by both neutralizing and non-nAbs.

## Results

### Characterization of new anti-gp350 mAbs

We generated and biochemically characterized new EBV gp350-specific mAbs, and evaluated their ability to neutralize EBV infection. In addition, we used the antibodies to map immunodominant epitopes on the EBV gp350 protein. To generate hybridomas, we immunized BALB/c mice with purified UV-inactivated EBV, boosted them with virus-like particles (VLPs) that incorporate the EBV gp350 ectodomain on the surface to enrich for production of anti-gp350 antibodies, then isolated splenocytes from the immunized mice and fused them with myeloma cells. We used indirect ELISA to screen supernatants from the hybridomas for specificity against purified EBV gp350 ectodomain protein (aa 4–863) and identified 23 hybridomas producing gp350-specific antibodies.

We determined the isotypes of the new antibodies to be IgG1 (n=14), IgG2a (n=5), IgG2b (n=1), a mixture of IgG1 and IgG2b (n=1), and a mixture of IgG1 and IgM (n=2). We found that all 23 hybridomas that produced antibodies (designated HB1-23) recognized the gp350 antigen in an initial ELISA screening using unfractionated and unpurified hybridoma supernatants (data not shown). We used affinity purification with protein A followed by SDS-PAGE to confirm the purity of all 23 antibodies. When we re-evaluated quantified amount of the purified antibodies (10 μg/ml) using indirect ELISA, all of the 23 antibodies had ELISA signals two times greater than those of phosphate buffered saline (negative control), and were considered as positive or specific to gp350. Of these, five (HB4, HB5, HB7, HB13, and HB14) demonstrated binding affinity equal to or greater than that of the positive control, nAb-72A1 (Fig. 1A). This difference in binding of the 23 antibodies could be due to differential exposure of cognate epitopes on gp350 in the assay performed.

**FIG 1.**
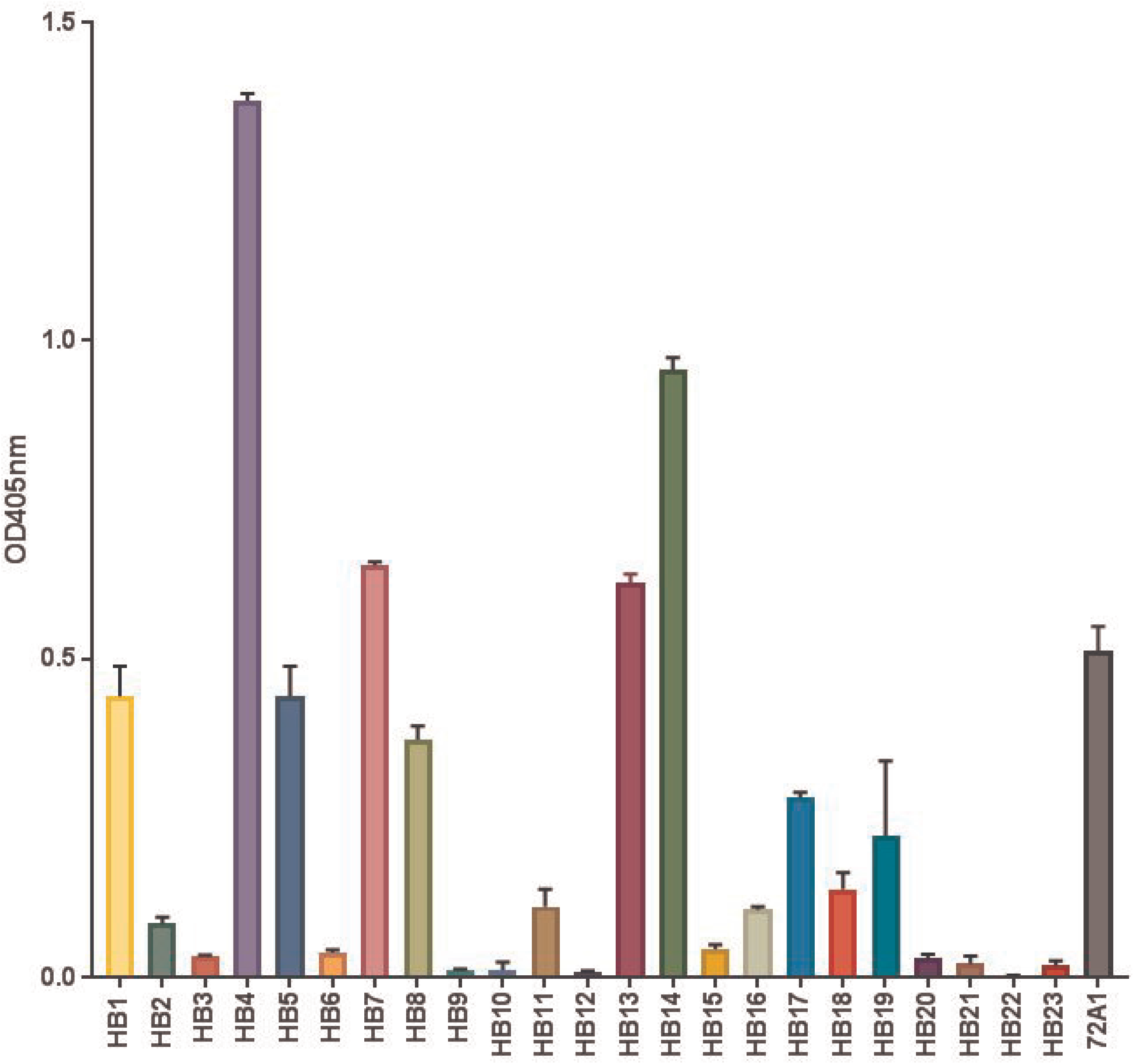
Specificity of anti-gp350 antibodies. (A) ELISA screening of hybridoma (HB) supernatants for anti-gp350-specific antibodies. Soluble EBV gp350 protein was used as the target antigen at 0.5 μg/ml. nAb-72A1 at 10 μg/ml and 1x phosphate buffered saline (PBS) were used as positive and negative (not shown) controls, respectively. Bound antibodies were detected using HRP-conjugated anti-mouse IgG (1:2,000). Twenty-three HB clones with ELISA signals two times greater than those of PBS control were considered as positive hybridomas. (B) Determining specificity of anti-gp350-producing hybridoma supernatants by immunoblotting with gp350-transfected stable CHO lysate. (C) Flow cytometric analysis of surface expression of gp350 protein on gp350-expressing CHO cells. Cells were stained with anti-gp350 mAb (1:250), followed by secondary goat anti-mouse conjugated to AF488.

Determining the nature of the binding between an antibody and its target antigen is an important consideration for the performance and specificity of an antibody, as it can involve the recognition of a linear or conformational epitope (26). We characterized the antibodies using immunoblot analysis of denatured gp350 antigen expressed from Chinese hamster ovary (CHO) cells, and showed that 16 of the antibodies reacted to both the 350 kDa and the 220 kDa splice variant. In contrast, HB2, HB3, HB6, HB7, HB13, HB20, and HB21 failed to recognize either of the denatured isoforms of gp350 (Fig. 1B). We further characterized the antibodies using flow cytometric analysis of CHO cells stably expressing gp350 on the cell surface, and revealed that HB1, HB2, HB3, HB5, HB6, HB9, HB11, HB12, HB15, HB17, HB19, HB20, and HB21 antibodies readily recognized gp350 (Fig. 1C). Given that HB2, HB3, HB20, and HB21 detected gp350 by flow cytometry, but not by immunoblot, suggests that these four antibodies recognized conformational epitopes (native) on gp350, whereas HB5, HB9, HB11, HB15, HB17, and HB19 recognized both linear and conformational epitopes (Fig. 1B-C). The observation that all 23 anti-gp350 antibodies recognized the gp350 antigen either by indirect ELISA, flow cytometry, or immunoblot assay suggests that we successfully produced antibodies that are specific to EBV gp350 protein.

### Analysis of the variable heavy and variable light chain sequences

We determined the sequences of the heavy and light chain variable region genes (V_H_ and V_L_, respectively) of the 23 new anti-gp350 antibodies, as well as nAb-72A1, and compared the sequences to published nAb-72A1 sequences (10, 33). The sequence of the CDR of this antibody was recently determined and published, revealing two unique IgG1 heavy chains and two unique light chains, one kappa and one lambda (10, 33). We used PCR to amplify the genes encoding the Vh and V_L_ chain regions in cDNA generated from the 23 hybridoma cells, as well as from HB168 (nAb-72A1). The PCR products presented distinct bands at approximately 350–400 bp and 450–500 bp for V_H_ and V_L_, respectively (data not shown). We sequenced purified fragments using Illumina MiSeq, followed by *in silico* analysis and identified CDRs for both V_H_ and V_L_ (Fig. 2). We identified two V_H_ and V_L_ sequences of nAb-72A1 as >94% identical to the previously published sequences (10), suggesting that nAb-72A1 exists as a mixed antibody, instead of the reported mAb (33). Similar to nAb-72A1, HB4, HB13, HB15, and HB23 hybridomas each produced a mixture of two antibodies, with unique sequences of the V_H_ chain showing at >5% frequencies, suggesting that they are not mAbs (Table 1). We were unable to identify coding sequences for V_L_ chains for HB7, HB9, and HB17, unless the frequencies were lowered to >1% (Table 1); in this case, the identified coding V_L_ chain sequences were identical.

**FIG 2.**
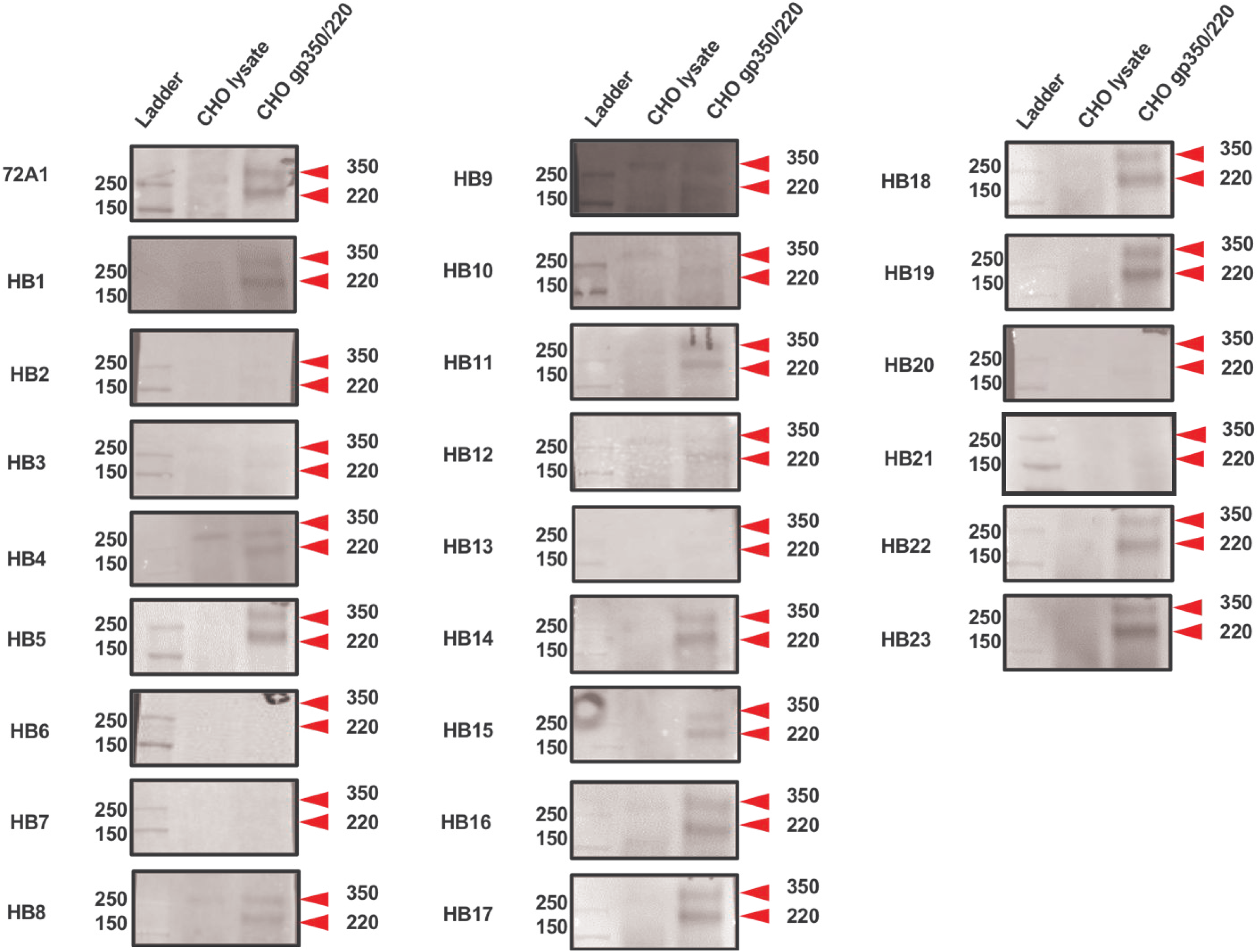
PROMALS3D multiple sequence alignment of (A) V_H_ and (B) V_L_ regions of 15 mAbs and nAb-72A1. The highly variable complementarity determining regions (CDR) 1–3, indicated by black boxes, define the antigen binding specificity. The conserved framework regions (FR) 1–4 flank the CDRs. Consensus amino acids (aa) are in bold and upper case. Consensus-predicted secondary structure (ss) symbols: alpha-helix, h; beta-strand, e.

**Table 1.**
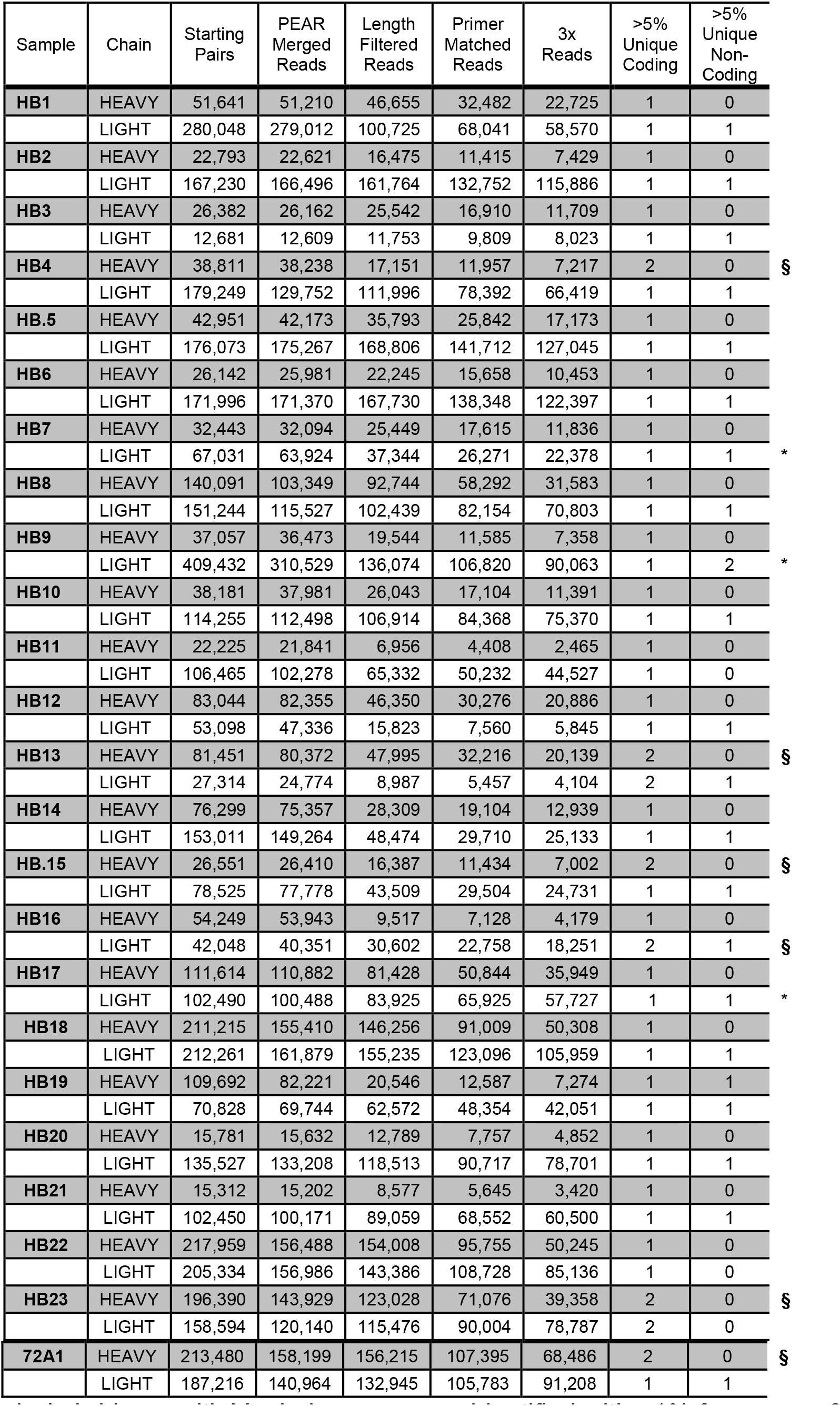
Summary of Illumina Dual Demultiplex of V_H_ and V_L_ regions.

* = hybridoma with V_L_ chain sequences identified with >1% frequency, § = hybridoma with more than one unique, plausible-coding V_H_ chain sequence with > 5% frequency. The term “unique” refers to unique sequence counts (so, identical sequences found in a substantial frequency of merged reads, not necessarily unique compared to other samples).

Our analysis and comparison of the V_H_ and V_L_ chain gene sequences of the 23 hybridomas compared to HB168 (nAb-72A1) showed unique sequences within the CDR 1–3 regions. Only HB8 and HB18 had identical V_H_ and V_L_ chain gene sequences, suggesting that the two are the same clone isolated separately; therefore, HB18 was excluded from subsequent experiments. One of the two HB15 antibodies had identical V_H_ and V_L_ gene sequences to that of HB10; however, based on the previous characterization, the presence of the additional antibody in HB15 was sufficient to confer subtle differences in biochemical characterizations for gp350 between the two antibodies. Thus, sequence analysis (Fig. 2) demonstrated that we generated 15 unique anti-gp350 mAbs, with distinct biochemical properties and sequence identities from the commercially available nAb-72A1.

### Neutralization assay

We evaluated the ability of the 15 mAbs (10 μg/ml or 50 μg/ml) to neutralize purified eGFP-tagged AGS-EBV infection of the Raji B cell line *in vitro* following standardized procedures (25) and determined the percentage of eGFP+ cells using flow cytometry as described (17, 18). We used the nAbs 72A1 and anti-gH/gL (E1D1) as positive controls, whereas the non-neutralizing mAb 2L10 was used as a negative control. Because HB4, HB7, HB13, HB15, HB16, HB19, HB21, and HB23 were confirmed to be mixtures based on isotyping or sequence data, we eliminated them from further consideration in the neutralization assay. We considered an antibody to be a neutralizer if it inhibited EBV infection >20% at 10 μg/ml and >60% at 50 μg/ml. Several mAbs inhibited EBV infection in a dose-dependent manner. HB20 and HB17 were the most effective in neutralizing EBV infection of Raji cells *in vitro*, whereby they reduced infection by 40% and >60%, and >30% and 80%, at 10 μg/ml and 50 μg/ml, respectively (Fig. 3). The HB9 and HB10 antibodies prevented EBV infection of Raji cells by ~25% at 10 μg/ml and ~60% at 50 μg/ml. The HB11 antibody neutralized <20% at 10 μg/ml, but showed a dose-dependent increase at 50 μg/ml by neutralizing EBV infection by 60%. By the set neutralization parameters, HB1-3, HB5-8, HB12, HB14, and HB22 did not neutralize EBV infection of Raji cells. In comparison, both nAb-72A1 and nAb-E1D1 neutralized EBV infection by >70% and 40%, respectively, at 10 μg/ml. The nAb-72A1 neutralized 100% of EBV infection at 50 μg/ml, whereas nAb-E1D1 neutralized infection by >60% at 50 μg/ml. As expected, the negative control, mAb-2L10, did not neutralize viral infection at either concentration. From the neutralization assay results, we organized the remaining 15 mAbs into two distinct groups, neutralizers (+) and non-neutralizers (-) as summarized in Table 2.

**FIG 3.**
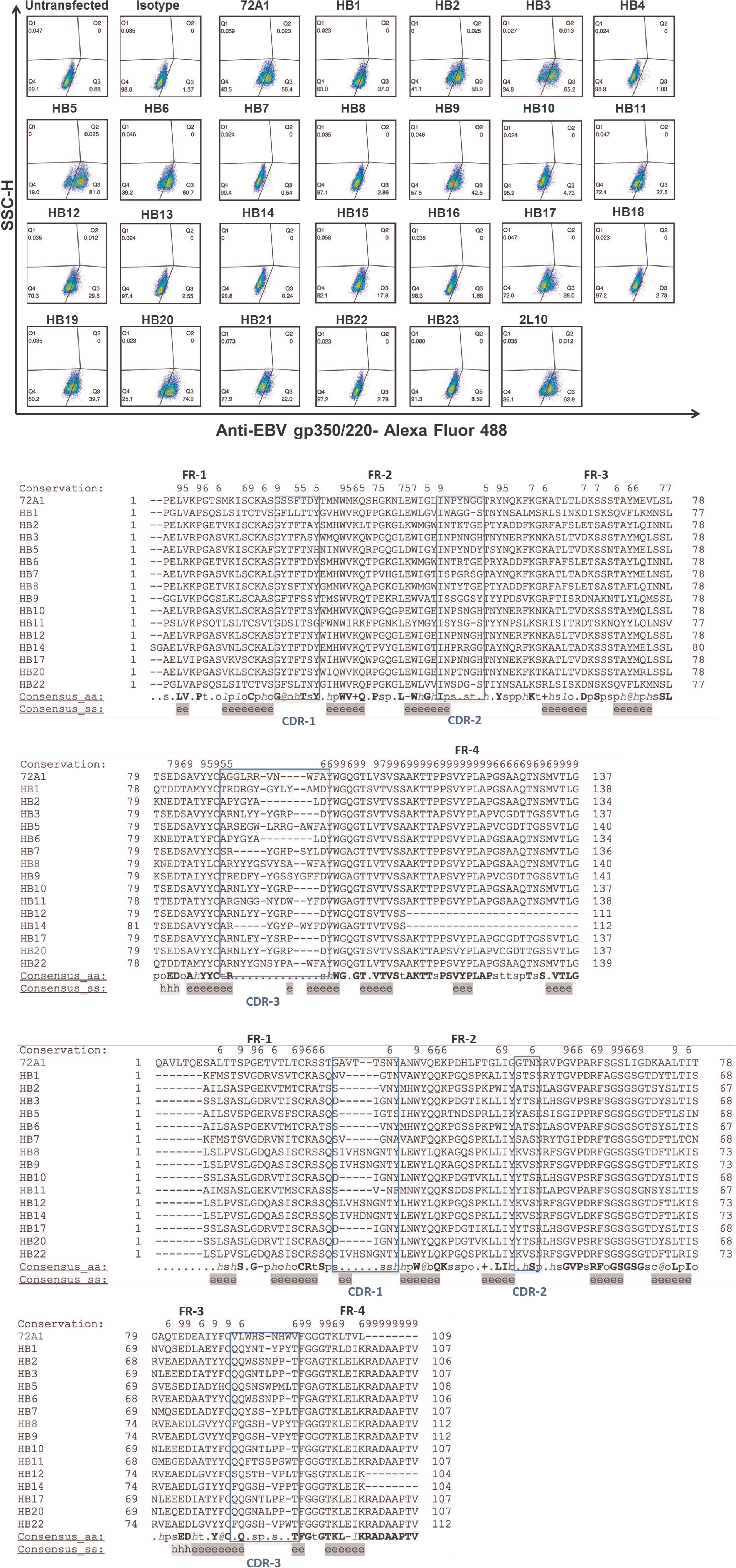
Neutralization activity of novel anti-gp350 mAbs against EBV-eGFP in Raji cells. EBV-eGFP was preincubated with 15 anti-gp350 mAbs at (A) 10 μg/ml and (B) 50 μg/ml, followed by incubation with Raji cells for 48 h. EBV-eGFP+ cells were enumerated using flow cytometry. Anti-gp350 (nAb-72A1) and anti-gH/gL (E1D1) mAbs served as positive controls and non-neutralizing anti-gp350 (2L10) mAb served as negative control.

**Table 2:** Summarized biochemical and functional characterizations of anti-gp350 antibodies.

| Antibody | IgG sub-class | Light chain | ELISA binding to purified EBV gp350/220 | FACS (CHO Cells) | Western blot | Neutralization activity |
| --- | --- | --- | --- | --- | --- | --- |
| HB1 | IgG1 | K | + | + | + | - |
| HB2 | IgG2a | K | + | + | - | - |
| HB3 | IgG2a | K | + | + | - | - |
| HB4 | IgG1 | K | + | - | + | ND |
| HB5 | IgG2a | K | + | + | + | - |
| HB6 | IgG1 | K | + | + | - | - |
| HB7 | IgG1 | K | + | - | - | - |
| HB8 | IgG1 | K | + | - | + | - |
| HB9 | IgG2a | K | + | + | + | + |
| HB10 | IgG1 | K | + | - | + | + |
| HB11 | IgG1 | K | + | + | + | + |
| HB12 | IgG1 | K | + | + | + | - |
| HB13 | IgG1 | K | + | - | - | ND |
| HB14 | IgG1 | K | + | - | + | - |
| HB15 | IgG1 | K | + | + | + | ND |
| HB16 | IgG1/ IgM | K | + | - | + | ND |
| HB17 | IgG2b | K | + | + | + | + |
| HB19 | IgG1/ IgM | K | + | + | + | ND |
| HB20 | IgG2a | K | + | + | - | + |
| HB21 | IgG1/IgG2b | K | + | + | - | ND |
| HB22 | IgG1 | K | + | - | + | - |
| HB23 | IgG1 | K | + | + | + | ND |
| 72A1 | IgG1 | K | + | + | + | + |
| E1D1<br>(anti gH/gL) | IgG1 | K | - | - | - | + |

ND=Not determined, + = positive, - = negative, ELISA = enzyme-linked immunosorbent assay, k = Kappa

### Epitope Mapping

We used the 15 new anti-gp350 mAbs (neutralizers vs. non-neutralizers) to identify most, if not all, of the relevant immunodominant aa residues targeted by both the neutralizing and non-nAbs. We used a RepliTope approach, in which overlapping peptides [15-mer with 11 aa overlap] that cover the complete sequence of gp350 (aa 1–907) were immobilized on microarray slides and probed with the purified anti-gp350 antibodies in an ELISA format. The nAb-72A1 was as positive control, because the cognate epitopes bound by the antibody have previously been reported.

We showed that two epitopes, _253_**TPIPGTGYAYSLRLTPRPVSRFL**_275_ and _393_**VSGLGTAPKTLIITRTATNATTT**_415_, were both bound by all 15 mAbs, as well as nAb-72A1 and 2L10, regardless of their neutralizing or non-neutralizing capabilities (Fig. 4). This consensus suggests that these epitopes are immunodominant. Several of the 15 mAbs (HB2, HB3, HB8, HB11, HB12, HB14, HB17, and HB22), as well as nAb-72A1, bound to _341_**ANSPNVTVTAFWAWPNNTE**_359_. Two epitopes, aa 341–359 and aa 393–415, were found within the previously identified single epitope II, which is encoded by nucleotides between 3,186 bp and 3,528 bp, corresponding to aa 326–439 of gp350 (37). Two mAbs, HB1 and HB10, bound _605_**TTPTPNATGPTVGETSPQA**_623_, an epitope located within the gp350 (aa 501–699) splice region that is involved in generation of the 220 kDa splice variant. A total of eight mAbs (HB1-3, HB8, HB10-12, and HB22) also bound to the region between _1_**MEAALLVCQYTIQSLIHLTGEDPG**_24_, which includes a region homologous to C3d, another molecule known to interact with CD21 (15, 33). Two epitopes common between most neutralizing and non-nAbs, _821_**PPSTSSKLRPRWTFTSPPV**_839_ and _875_**LLLLVMADCAFRRNLSTSHTYTTPPY**_899_, were located upstream and downstream, respectively, of the transmembrane domain on the C-terminus of gp350. Epitope aa 821–839 is located within the previously identified epitope I, which is located between aa 733–841 (37). Furthermore, epitope aa 821–839 is potentially involved in EBV infection of B cells (35). However, our study could not identify two epitopes located at aa 282–301 and aa 194–211, which were previously shown to be involved in the binding of nAb-72A1 and CD21, respectively (30, 33, 35). Similar to previously reported data, we showed that nAb-72A1 bound _145_**EMQNPVYLIPETVPYIKWDN**_164_, one of the neutralizing epitopes on gp350 (30, 33, 35).

**FIG 4.**
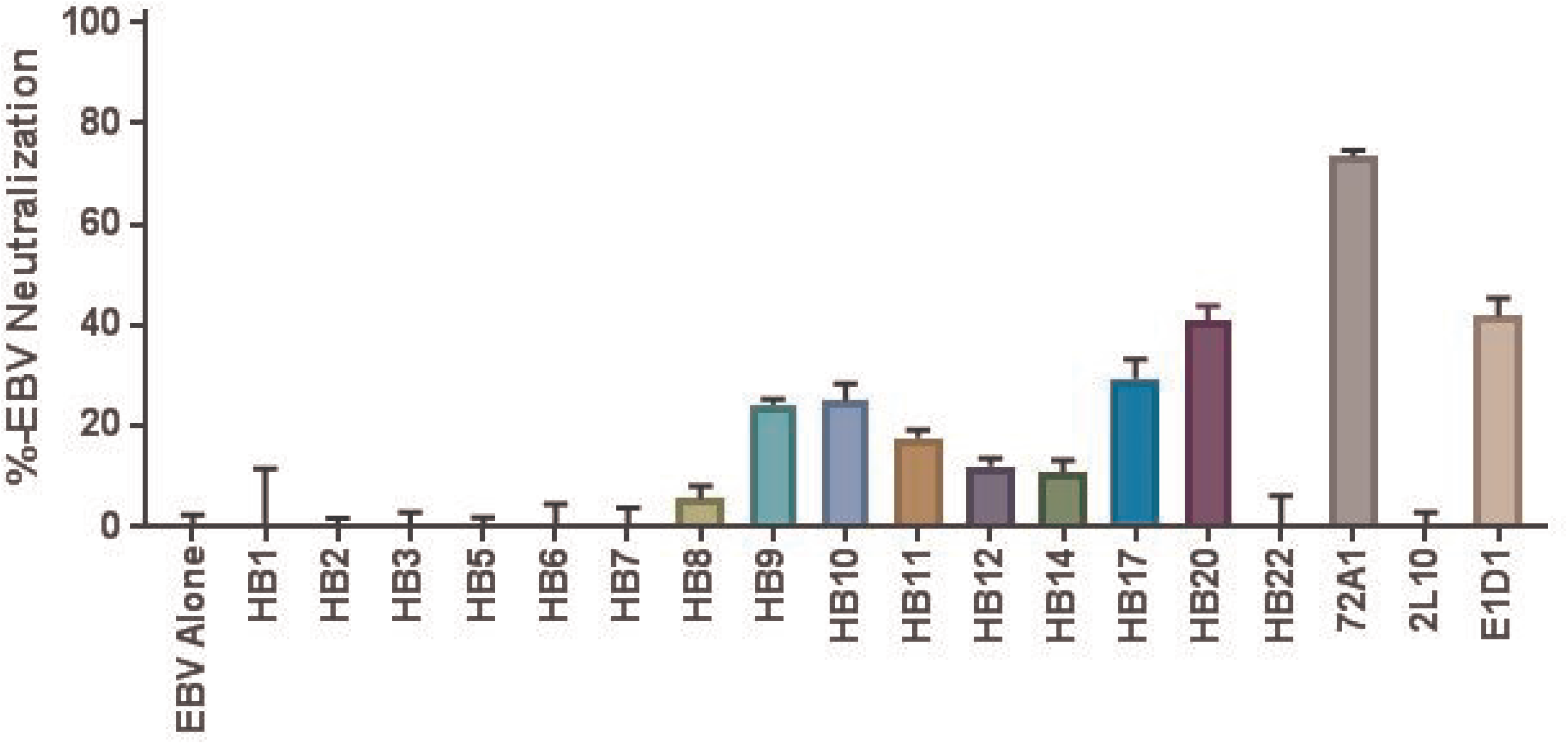
Identification of nine gp350 epitopes using 15 neutralizing and non-neutralizing mAbs. Residues in bold represent the gp220 splice variant region. Residues in red represent RepliTope-identified epitopes and exact residues. Italic residues represent canonical neutralizing epitope, underlined residues represents epitope bound by all assayed mAb.

Analysis of V_H_-V_L_ sequences from the HB168 (nAb-72A1) hybridoma by our group and others revealed that the hybridoma produces two antibodies: one that is gp350-specific and another that recognizes mineral oil-induced plasmacytoma (MOPC)(10). To further interrogate gp350 for additional neutralizing epitopes, we used the gp350-specific nAb-72A1 V_H_-V_L_ sequence to generate chimeric (mouse/human) recombinant antibodies. Similarly, we used the V_H_-V_L_ sequence for the HB20 antibody, which our neutralization analysis above showed to be one of the most effective nAbs, to generate chimeric antibodies. We generated a negative control chimeric recombinant antibody using V_H_-V_L_ sequences from the gp350-specific but non-neutralizing HB5 antibody described above (Fig. 3).

We performed comparative analysis of the epitope binding pattern using the chimeric recombinant antibodies and revealed a novel epitope, _381_**GAFASNRTFDIT**_392_, which was bound by HB20 and nAb-72A1, but not by HB5, and is distinct from the _145_**EMQNPVYLIPETVPYIKWDN**_164_ epitope, which is bound only by the nAb-72A1 (Fig. 5).

**FIG 5.**
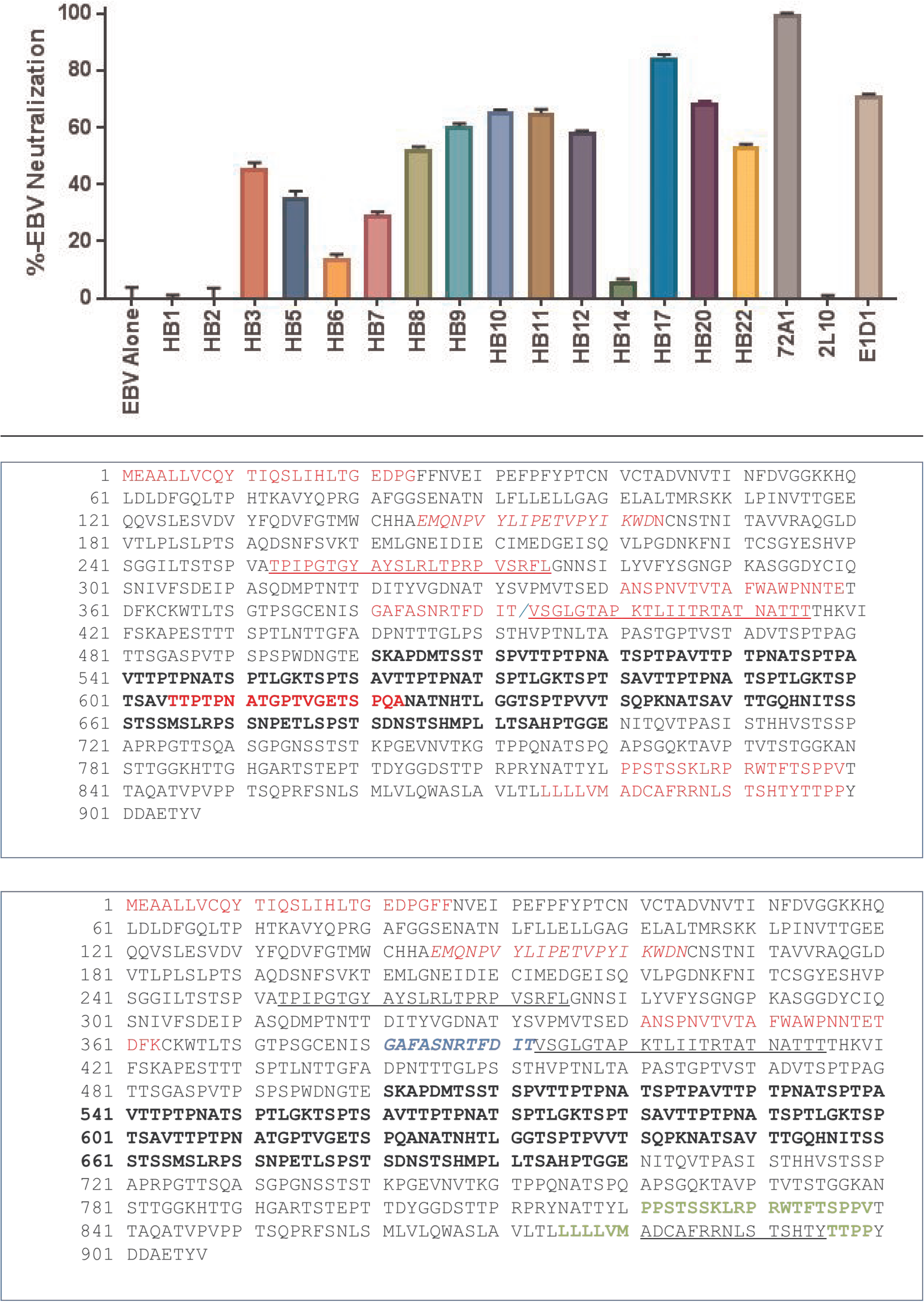
Identification of novel gp350-neutralizing epitope by epitope mapping of neutralizing (nAb-72A1 and HB20) vs. non-neutralizing (HB5) anti-gp350 mAbs. Residue positions of nAb-72A1 (red), HB20 (blue), and HB5 (green) are indicated. Bold black residues represent splice variant region, underlined residues represent epitopes bound by nAb-72A1, HB20, and HB5. Italic residues represent epitopes bound by nAbs. Red residues represent epitopes bound by nAbs-72A1, blue residues represent residues bound by HB20 and green represents residues bound by HB5.

## Discussion

Despite more than 50 years of EBV vaccine research, few candidates have demonstrated even partial clinical efficacy, and none have been efficacious enough to elicit sterilizing immunity and be licensed (3). Antibodies, whether elicited in the host naturally or via passive immunization, provide an effective first-line of defense against viral infection. Several studies have indicated that the major immunodominant glycoprotein EBV gp350 is an ideal target for EBV nAbs production. Although the ectodomain of EBV gp350 (aa 1–841) contains at least eight unique CD21 binding epitopes, only one of these epitopes (aa 142–161) is capable of eliciting nAbs (30, 32, 35). Although the aa residues that constitute the other epitopes and their role in generating nAbs have not been elucidated, this information would be valuable in the precise design of effective EBV peptide vaccines. To date, nAb-72A1 remains the only EBV antibody with proven clinical prophylactic efficacy; it confers shortterm protection by reducing and delaying EBV infection onset in immunized pediatric transplant patients (21).

In this study we generated 23 hybridomas producing gp350-specific antibodies and characterized their ability to bind gp350 protein (Fig. 1). Out of the 23 hybridomas, we determined that 15 were monoclonal and novel based on their V_H_ and V_L_ CDR sequences, compared to the reported sequence of nAb-72A1 (22) (Fig. 2)

Following confirmation that the new mAbs recognized EBV gp350 antigen and contained unique V_H_-V_L_ sequences, our further characterization revealed that mAbs HB9, HB10, HB11, HB17, and HB20 inhibited EBV infection in a dose-dependent manner, with HB17 and HB20 being the best neutralizers (Fig. 3). Thus, our study provides new neutralizing and non-nAbs against EBV infection, which could potentially inform future EBV vaccine research.

Various methods, including lectin/ricin immune-affinity assay (34), purified mAbs (1, 22), purified soluble gp350 mutants, synthetic peptides (15, 16, 33), cell binding assays (35), and crystal structure of partial gp350 protein (aa 4–443) (30), have been used to identify the critical gp350 epitopes responsible for its interaction with the CD35 and CD21 cellular receptors (*for a detailed summary review* see Table 3). We used an epitope mapping assay to identify a total of nine epitopes, including two new epitopes, and one bound by all mAbs, including nAb-72A1 and 2L10, regardless of the mAb neutralizing capability to EBV infection. This suggests it is an immunodominant epitope on the gp350 protein. We further identified portions of four of the previously described epitopes, including the only currently recognized neutralizing epitope, and defined their exact aa residues. However, we could not identify two previously reported epitopes located at aa 282–301 and aa 194–211, which have been reported to be involved in the binding of nAb-72A1 or CD21, respectively (30, 33, 35) (Fig. 4).

**Table 3.** Summary of gp350 epitope mapping over time using various methodologies.

| Method | mAbs/protein/ peptides | Number of epitopes identified | Reference |
| --- | --- | --- | --- |
| Competitive binding assay: Tagged mAb vs. untagged mAb | Newly generated mAbs | 8 epitopes – Sequence not defined | Qualtiere et al., 1987 (21) |
| Binding studies: Determine the effects of anti-gp350 mAbs on gp350 binding to CR2 | Newly generated mAbs | 2 possible regions identified by sequence alignment to C3d sequence:<br>1. aa 21–28<br>2. aa 372–378 | Nemerow et al., 1987 (16) |
| Peptide digest and immunoprecipitation | Truncated and mutant protein; mAbs (72A1 and BOS-1) | Narrowed down to the first 470 residues | Tanner et al., 1988 (32) |
| Binding studies | Peptide and protein | 2 sequences defined:<br>1. aa 21–28<br>2. N-terminus of gp350 | Nemerow et al., 1989 (15) |
| Dot Blot immunoassay: Purified truncated protein incubated with mAbs | Protein – 8 clones overlapping N- and C-terminal portions of protein; mAbs from Qualtiere *et al.*, 1987 | 3 sequences defined:<br>1. aa 310–325<br>2. aa 326–439<br>3. aa 733–841 | Zhang et al., 1991 (37) |
| Peptide cell binding assay to 2 CR2-positive (Raji and Ramos) and 1 -negative (P3HR-1) cell lines | Synthesized peptides covering gp350 (907 aa) | 7 regions, 3 identified:<br>1. aa 142–161<br>2. aa 282–301<br>3. aa 822–841 | Urquiza et al., 2005 (35) |
| Crystal structure and binding studies | Mutant proteins; mAbs 72A1 | 3 epitopes (based on 72A1 binding and gp350 4-443)<br>1. aa 16-29<br>2. aa 142-161<br>3. aa 282-301 | Szakonyi et al., 2006 (30) |
| Structural docking studies and antigenicity mapping | gp350 and CR2 crystal structure alignment/docking | Single epitope (based on gp350 aa 1–470)<br>1. aa147–165 | Sitompul et al., 2012 (27) |
| Structural alignment:<br>computer modeling of<br>gp350 and 72A1 and<br>docking studies | Peptides (used in<br>immunization); mAb (72A1) | 4 epitopes: identified<br>1. aa 14-20<br>2. aa 144-161<br>3. aa 194-211<br>4. aa 288-291 | Tanner et al., 2015<br>(33) |

Our comparative epitope mapping analysis results identified a novel neutralizing epitope, _381_**GAFASNRTFDIT**_392_, which was bound preferentially by the nAbs HB20 and nAb-72A1, but not the nonneutralizing mAb HB5 (Fig. 5). The new neutralizing epitope is distinct from the reported canonical nAb-72A1-binding epitope aa 145–164, on gp350, suggesting that epitope 381–392 is a novel epitope on gp350 that might be capable of eliciting nAbs.

In conclusion, in this study we generated 15 novel EBV gp350-specific mAbs, characterized their binding to gp350, determined their neutralization activity against EBV infection *in vitro*, mapped their cognate epitopes, and defined the exact aa residues they recognize on gp350. We confirmed six of the eight previously described epitopes responsible for generating neutralizing and non-nAbs and defined the exact aa residues they bind. This study also confirmed that the binding epitopes on gp350 that elicit nAbs are between aa 4–443 (30). We identified an additional neutralizing epitope and two new non-neutralizing epitopes, with one located downstream of the gp350 ectodomain (aa 1–841). The newly developed mAbs will be useful research tools for informing future vaccine development, diagnosis of viral infection, or therapeutic/prophylactic management of post-transplant lymphoproliferative diseases, either individually, in combination with nAb-72A1, or with other mAbs such as neutralizing anti-gH/gL (E1D1).

## Materials and Methods

### Cells and viruses

EBV-AGS, a human gastric carcinoma cell line infected with a recombinant Akata virus expressing enhanced fluorescent green protein (eGFP) was a kind gift of Dr. Liisa Selin (University of Massachusetts Medical School). Anti-EBV gH/gL (E1D1) hybridoma cell line was a kind gift of Dr. Lindsey Hutt-Fletcher (Louisiana State University Health Sciences Center). Chinese hamster ovary cells (CHO); human embryonic kidney cells expressing SV-40 T antigen (HEK-293T); HEK-293 6E suspension cells; EBV-positive Burkitt lymphoma cells (Raji); myeloma cells (P3X63Ag8.653); and anti-EBV gp350 nAb-72A1 hybridoma cells (HB168) were purchased from American Type Culture Collection (ATCC). EBV-AGS cells were maintained in Ham’s F-12 media supplemented with 500 μg/ml neomycin (G418, Gibco). Raji, P3X63Ag8.653, and HB168 hybridoma cells were maintained in RPMI 1640. CHO and HEK-293T cells were maintained in DMEM. HEK-293 6E cells were maintained in FreeStyle F17 Expression Medium supplemented with 0.1% Pluronic F-68. All culture media were supplemented with 10% fetal bovine serum (FBS), 2% penicillin-streptomycin, and 1% l-glutamine, with the exception of Freestyle F17 expression medium.

### Antibodies and plasmids

Primary antibodies: EBV anti-gp350 nAb (72A1) and anti-gH/gL (E1D1) were purified from the supernatant of HB168 and E1D1 hybridoma cell lines, respectively using Capturem™ Protein A Maxiprep spin columns (Takara). Anti-gp350/220 mAb (2L10) was purchased from Millipore Sigma.

Secondary antibodies: Horseradish peroxidase (HRP)-conjugated goat anti-mouse IgG for immunoblot or ELISA were purchased from Bio-Rad. Alexa Fluor® (AF) 488-conjugated goat anti-mouse IgG (H+L) for flow cytometry was purchased from Life Sciences Tech. Goat anti-mouse IgG (H+L) secondary antibody and DyLight 650 for epitope mapping were purchased from Thermo Fisher Scientific.

The construction of the pCI-puro vector and pCAGGS-gp350/220-F has been described (17, 20).

### Virus production and purification

eGFP-tagged EBV was produced from the EBV-infected AGS cell line as described (29). Briefly, EBV-AGS cells were seeded to 90% confluency in T-75 flasks in Ham’s F-12 medium containing G418 antibiotic. After the cells reached confluency, G418 media was replaced with Ham’s F-12 medium containing 33 ng/ml 12-O-tetradecanoylphorbol-13-acetate (TPA) and 3 mM sodium butyrate (NaB) to induce lytic replication of the virus. Twenty-four h post-induction, the media was replaced with complete Ham’s F-12 media without G418, TPA, or NaB and cells were incubated for 4 days at 37°C. The cell supernatant was collected, centrifuged, and filtered using a 0.8-μm filter to remove cell debris. The filtered supernatant was ultra-centrifuged using a Beckman-Coulter type 19 rotor for 70 min at 10,000 rpm to pellet the virus. EBV-eGFP virus was titrated in both HEK-293T cells and Raji cells, and stocks were stored at −80°C for subsequent experiments.

### Generation and purification of gp350 virus-like particles

To generate gp350 VLPs, equal amounts (8 μg/plasmid) of the relevant plasmids (pCAGGS-Newcastle disease virus [NDV]-M, -NP, and gp350 ectodomain fused to fusion protein cytoplasmic and transmembrane domains) were co-transfected into 80% confluent CHO cells seeded in T-175cm^2^ flasks; supernatant from transfected cells was collected and VLPs were purified and composition characterized as previously described (18).

### Production of hybridoma cell lines

Seven days prior to immunization, two eight-week-old BALB/c mice were bled for collection of pre-immune serum. The mice were immunized with purified UV-inactivated EBV three times (Day 0, 21, and 35) and boosted every 7 days three times (Day 42, 49, and 56) with VLPs incorporating gp350 on the surface after Day 35. The mice were sacrificed and their splenocytes were isolated, purified, and used to fuse with P3X63Ag8.653 myeloma cells at a ratio of 3:1 in the presence of polyethylene glycol (PEG, Sigma). Hybridoma cells were seeded in flat-bottom 96-well plates and selected in specialized hybridoma growth media with HAT (Sigma) and 10% FBS.

### Indirect ELISA

Hybridoma cell culture supernatant from wells that had colony-forming cells were tested for antibody production by indirect ELISA. Briefly, immunoplates (Costar 3590; Corning Incorporated) were coated with 50 μl of 0.5 μg/ml recombinant EBV gp350/220 (Millipore) protein diluted in phosphate buffered saline (PBS, pH 7.4) and incubated overnight at 4°C. After washing three times with PBS containing 0.05% (v/v) Tween 20 (washing buffer), plates were blocked with 100 μl washing buffer containing 2% (w/v) bovine serum albumin (BSA), incubated for 1 h at room temperature, and washed as above. 100 μl of hybridoma supernatant was added to each well (in triplicate) and incubated for 2 h at room temperature. PBS and nAb-72A1 were added as negative and positive controls, respectively. The plates were washed as described above, followed by incubation with goat anti-mouse IgG horseradish peroxidase-conjugated secondary antibody (1:2,000 diluted in PBS) at room temperature for 1 h. The plates were washed again and the chromogenic substrate 2,2’-azino-bis (3-ethylbenzothiazoline-6-sulphonic acid) (ABTS, Life Science Technologies) was added. The reaction was stopped using ABTS peroxidase stop solution containing 5% sodium dodecyl sulfate (SDS) in water. The absorbance was read at an optical density of 405 nm using an ELISA reader (Molecular Devices). Three independent experiments were performed.

### Antibody purification, quantification, and isotyping

Hybridoma cells from selected individual positive clones were expanded stepwise from 96-well plates to T-75 flasks. At confluence in T-75 flasks, supernatant from individual clones was collected, clarified by centrifugation (4,000 g, 10 min, 4°C), and filtered through a 0.22-μm-membrane filter (Millipore). Antibodies were further purified using Capturem™ Protein A Maxiprep (Takara) and stored in PBS (pH 7.4) at 4°C. Antibodies were analyzed by SDS-PAGE to determine purity. Bicinchoninic acid assay (BCA assay; Thermo Fisher Scientific) was conducted to determine the concentration of purified antibodies. Isotype identification was performed with the Rapid ELISA mouse mAb isotyping kit (Thermo Fisher Scientific).

### RNA extraction, cDNA synthesis, and sequencing of the variable region of the mAbs

Total RNA was extracted from 1×10^6^ hybridoma cells using the RNeasy Mini Kit (Qiagen). Each hybridoma clone cDNA was synthesized in a total volume of 20 μl using Tetro Reverse Transcriptase (200 u), RiboSafe RNase Inhibitor, Oligo(dT)18 primer, dNTP mix (10 mM each nucleotide), and 100–200 ng RNA. Reverse transcription was performed at 45°C for 30 min, and terminated at 85°C for 5 min. The cDNA was stored at −20°C. Immunoglobulin (Ig) V_H_ and V_L_ were amplified using the mouse Ig-specific primer set purchased from Novagen (12). The V_H_ and V_L_ genes were amplified in separate reactions and PCR products were visualized on 1% agarose gel.

The V_H_ and V_L_ amplicons were sequenced using an Illumina MiSeq platform: duplicate 50 μl PCR reactions were performed, each containing 50 ng of purified cDNA, 0.2 mM dNTPs, 1.5 mM MgCl_2_, 1.25 U Platinum Taq DNA polymerase, 2.5 μl of 10x PCR buffer, and 0.5 μM of each primer designed to amplify the V_H_ and V_L_. The amplicons were purified using AxyPrep Mag PCR Clean-up kit (Thermo Fisher Scientific). The Illumina primer PCR PE1.0 and index primers were used to allow multiplexing of samples. The library was quantified using ViiA^™^ 7 Real-Time PCR System (Life Technologies) and visualized for size validation on an Agilent 2100 Bioanalyzer (Agilent Technologies) using a high-sensitivity cDNA assay. The sequencing library pool was diluted to 4 nM and run on a MiSeq desktop sequencer (Illumina). The 600-cycle MiSeq Reagent Kit (Illumina) was used to run the 6 pM library with 20% PhiX (Illumina), and FASTQ files were used for data analysis(19).

### Chimeric mAb construct generation

To generate chimeric mAbs, the V_H_ and V_L_ sequences were cloned into the dual-vector system pFUSE CHIg/pFUSE CLIg (InvivoGen), which express the constant region of the heavy and light chains of human immunoglobulins, respectively (Genewiz). The constructs were transiently transfected into HEK-293 6E cells. The supernatants were collected at 72 h post-transfection and IgG was purified using protein A/G affinity chromatography.

### Immunoblot analysis

CHO cells were cultured and stably co-transfected with pCAGGS-gp350/220 F and pCI-puro vector containing a puromycin resistance gene. Forty-eight h post-transfection, DMEM media containing 10 μg/ml of puromycin was added to enrich for cells expressing gp350 protein. Puromycin-resistant clones were expanded, followed by flow cytometry sorting using nAb-72A1 to a purity >90%. EBV gp350-positive CHO cells were harvested and lysed in radioimmunoprecipitation assay buffer (RIPA) followed by centrifugation at 15,000 g for 15 min on a benchtop centrifuge. The supernatants were collected and heated at 95°C for 10 min in SDS sample buffer containing β-mercaptoethanol, then separated using SDS-PAGE. Proteins were transferred onto a nitrocellulose membrane using an iBlot™ Transfer System (Thermo Fisher Scientific) followed by incubation in blocking buffer (1% BSA; 20 mM Tris-HCl, pH 7.5; 137 mM NaCl; and 0. 1%Tween-20 [TBST]) for 1 h. The blots were incubated in TBST containing purified anti-gp350 antibodies (1:50) overnight at 4°C. After three washes with TBST, the blots were incubated with a goat anti-mouse conjugated to horseradish peroxidase (1:2,000) in TBST for 1 h. After three washes, the antibody-protein complexes were detected using the Amersham ECL Prime Western Blotting Detection Reagent (GE Healthcare). All experiments were independently repeated three times.

### Flow cytometry

To assess the ability of purified anti-gp350 mAb to detect surface expression of EBV gp350 protein by flow cytometry, CHO cells that stably express EBV gp350 were harvested and stained with purified anti-gp350 (10 μg/ml), followed by AF488 goat anti-mouse IgG secondary antibody. Flow cytometric analysis was performed on a C-6 FC (BD Biosciences) and data was analyzed using FlowJo Cytometry Analysis software (FlowJo, LLC) as described (18). All experiments were independently repeated three times.

### EBV neutralization assay

Purified individual anti-gp350 mAbs were incubated with purified AGS-EBV-eGFP (titer calculated to infect at least 20% of HEK293 cells seeded in 100 μl of serum-free DMEM) for 2 h at 37°C. To represent EBV infection of B cells, the pre-incubated anti-gp350 mAbs/AGS-EBV-eGFP were used to infect 5×10^5^ Raji cells seeded in a 96-well plate. Anti-gp350 neutralizing 72A1 and non-neutralizing 2L10 mAbs served as positive and negative controls, respectively. Plates were incubated at 37°C and the number of eGFP+ cells was determined using flow cytometry 48 h post-infection. All dilutions were performed in quintuplicate and the assays repeated three times for Raji cells. Antibody EBV neutralization activity was calculated as: % neutralization = (EBValone-EBVmAb) / (EBValone) x 100.

### Epitope mapping

Anti-gp350 mAbs were incubated with a multi-well EBV GP350/GP340 RepliTope (JPT) peptide microarray displaying 224 peptides (15-mers with 11 aa overlap) in 3×7 subarrays. Briefly, anti-gp350 mAbs were diluted in blocking solution (TBS-T and 2% BSA) to a final concentration of 10 μg/ml and incubated with the microarray slide for 1 h at 30°C with shaking. Slides were washed 5 times with wash buffer (TBS-T), followed by incubation with 1 μg/ml secondary antibody Dylight 650 (Thermo Fisher Scientific) for 1 h at 30°C. After washing 5 times with wash buffer and 2 times with distilled water, microarray slides were dried by centrifugation. Detection was performed using the Agilent DNA microarray scanner.

## Acknowledgments

We are grateful to Lindsey Hutt-Fletcher for providing the EBV gH/gL (E1D1) hybridoma cell line, Liisa Selin for providing EBV Akata virus expressing eGFP, and Chao Guo for technical assistance with the RepliTope protocol.

This work was supported by the National Institute of Health (NIH) grant R21CA205106 and a generous philanthropic donation from the V Foundation to J.G.O. Research reported in this publication also included work performed in the City of Hope Integrative Genomics Core, Flow Cytometry Core, and Animal Resource Center supported by the National Cancer Institute of the National Institutes of Health under award number P30CA033572. The content is solely the responsibility of the authors and does not necessarily represent the official views of the National Institutes of Health. We thank Dr. Sarah T. Wilkinson for editing of the manuscript and for her insightful feedback and discussion.

